# Pan-cancer scale landscape of simple somatic mutations

**DOI:** 10.1101/112367

**Authors:** Nan Zhou, Blanca Gallego, Jinku Bao, Guy Tsafnat

**Author notes:** To whom correspondence should be addressed Jinku Bao, Guy Tsafnat.

## Abstract

Genome is the carrier of somatic mutations during the development of cancer. The catalogue of simple somatic mutations (SSM) is a subgroup of somatic mutations. It includes single base substitutions, small deletions and insertions of <= 200 bp, and multiple base substitutions of <= 200 bp. The comprehensive landscape of SSM has not been studied. After analysed 46,692,922 SSM of 10,878 samples, we proposed a pan-cancer scale landscape of SSM for 60 cancer projects in ICGC. In addition, the whole genome sequencing (WGS) and whole exome sequencing (WXS) techniques were compared according to the landscape of SSM. The result indicates numbers of SSM vary dramatically in different cancers. WGS can detect 10 times more single base substitutions and insertions than WXS. In terms of WXS, it called 10 times more deletions than insertions. Multiple base substitutions have not been well studied so they were just observed in a few cancer projects. Cancers generally show high prevalence of C > T substitutions at NpCpG trinucleotide contexts. Skin cancer showed distinct mutational spectra. Breast cancer, bladder cancer, and cervical cancer were found to have similar mutational spectra. Acute myeloid leukemia and lung cancer from South Korea, and colorectal cancer from China show high density of single base substitutions per mega base in chromosome Y. To sum up, our study and findings will be thought provoking in studying SSM in cancer.

## Introduction

It is explicit that cancer genomes carry somatic mutations. Somatic mutations are generally categorised into substitutions, insertions and deletions (indels), copy number variations and structural variations [1,2]. Study of mutations of a single gene like p53 in cancer can date back to decades ago [3]. After the advent of high throughput sequencing technology around 2005, genome wide studies have unprecedentedly enriched data of somatic mutations in cancer. For example, projects like TCGA (The Cancer Genome Atlas) and ICGC (International Cancer Genome Consortium) have promoted sequencing of thousands of cancer genomes. In ICGC, somatic mutations are provided as simple somatic mutations (SSM). SSM consists of single base substitutions, deletions of <= 200 bp, insertions of <= 200bp, and multiple base substitutions (>= 2 bp and <= 200 bp), which form a subgroup of somatic mutations.

The characteristics of complex somatic mutations in cancers can be clearly shown by mutational landscape [4,5]. For instance, exogenous carcinogenic factors induced somatic mutations in lung cancer and melanoma have been identified by studying the landscape of mutations [6,7]. Nowadays, large scale landscape of somatic mutations of all main cancer types are also available from other studies [8,9]. They indicated that landscape of mutations is an insightful way to study somatic mutations in cancer.

Whole genome sequencing (WGS) and whole exome sequencing (WXS) are two widely used strategies in cancer genomics. The difference between somatic mutations generated by WGS and WXS has not been well studied. Landscape of SSM in ICGC has not been studied yet. In this study, we provided a comprehensive landscape of SSM of all available cancer projects in ICGC and compared the difference between WGS and WXS in generating SSM at the pancancer scale the first time.

## Materials and methods

### Data collection and preparation

SSM data of cancer genomes used in this study was from release 23 of ICGC [10,11]. It was released on December 7, 2016. It includes 70 cancer projects, 21 cancer primary sites, and 46,693,172 SSM. These data were downloaded according to projects. Because 10 projects lack SSM data, only SSM of 60 cancer projects were downloaded and analysed. Because a mutation can have multiple impacts in the genome, there are duplicated records in the data. Duplications are removed according to the ICGC mutation id. Mutations not found by WGS and WXS were also removed. Finally, the data used for further analysis contains 60 cancer projects, 10,878 samples and 46,692,922 unique SSM in total. Details about the data used in this study were available in Table S1.

### Chromosome size

The chromosome size of whole genome was from human reference genome UCSC hg19 [12]. The size of whole exome and length of exons in each chromosome were calculated from human exome of Ensemble release 75 [13]. The exome data was retrieved from Ensembl’s MySQL database using “select R.name, E.seq_region_start, E.seq_region_end, E.stable_id from exon as E,seq_region as R where E.seq_region_id=R.seq_region_id”. Length of exons in each chromosome was calculated subsequently. The chromosome size for WGS and WXS was listed in Table S2.

Number of mutations per mega base (Mb) in each chromosome for single base substitutions was calculated. In each chromosome, the total number of mutations was divided by the length in Mb of the corresponding chromosome, either using chromosome size of WGS or WXS.

### Mutational spectra

Single base substitutions were firstly classified into six types, C > A, C > G, C > T, T > A, T > C, T >G [14]. Each type was furthered divided into 16 trinucleotide spectra by integrating a flanking base immediate to the mutated base at 5’ and 3’. Finally, single base substitutions are categorised into 96 spectra. Indels and multi base substitutions were classified according to the length of mutated bases. Indels were categorised into four spectra, <= 3 bases, 4-10 bases, 11-30 bases, and >= 31 bases. Multi base substitutions were categorised into three spectra, 1-20 bases, 21-50 bases, and >= 51 bases.

### K-S test

In order to see whether a larger chromosome would carry more single base substitutions, Kolmogorov-Smirnov test (K-S test) was used to test if the distribution of mutations across chromosomes was drawn from the same distribution of chromosome length. Because the number of mutations and length of chromosomes are at different scale, they were normalized into range [0, 1] using feature scaling approach X’ = (X - X_min_) / (X_max_ - X_min_), where X is a numeric vector and X’ is the normalised X. The K-S test was implemented by R function ks.test, with parameter alternative set to two.sided [15]. A p value of >= 0.05 indicates two distributions are the same, otherwise significantly different.

## Results

### SSM in cancer

As can be seen in Figure 1, the number of single base substitutions dominates SMM. Deletions and insertions take a relative large portion while multiple base substitutions account for the smallest composition. Numbers of SSM found by different cancer projects vary dramatically. The median of single base substitutions varies from a few to more than 10,000. Compared to WXS, WGS is more likely to detect more insertions and multiple base substitutions. In each cancer project, the number of identified deletions is proportional to the number of insertions. That means if there are more deletions in a cancer, there should be more insertions, and vice versa.

**Figure 1.**
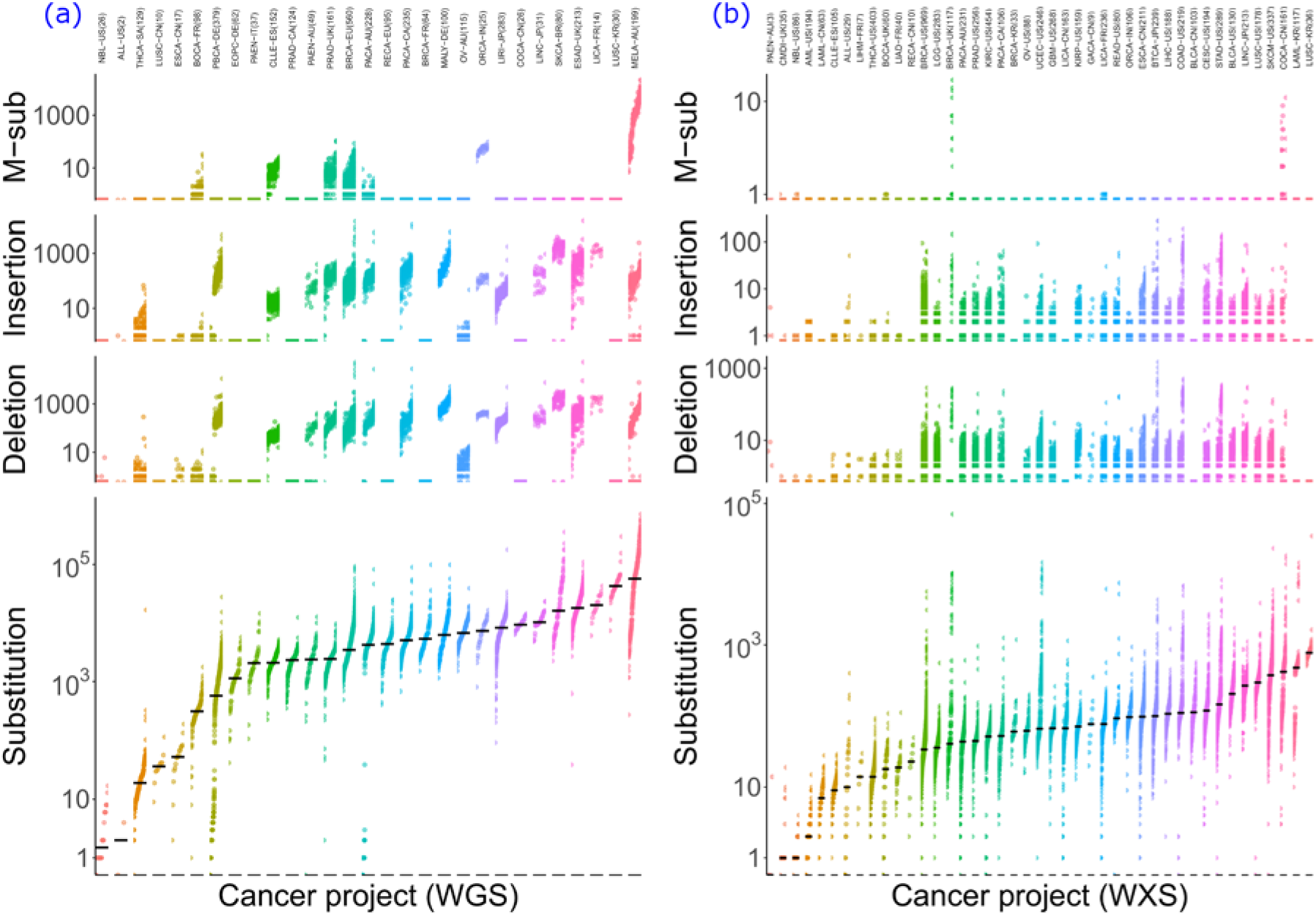
Distribution of SSM in all samples of 60 ICGC cancer projects. Each point denotes a sample. The number of mutations are on the y axis. For better perception, log10 scaling was used on y axis when creating the plot. Substitution explicitly denotes single base substitutions. M-sub represents multiple base substitutions. Black horizontal lines denote the median of mutations of all samples of each project. (a), Projects using WGS strategy. (b), Projects using WXS strategy. The colour code for cancer projects in (a) has no relation with that in (b), which is a routine in the context.

Regarding to WGS (Figure 1a), the majority of cancers have 1,000 to 100,000 single base substitutions. NBL-US contains approximately 10 single base substitutions, whereas at the opposite extreme, about 1/3 of MELA-AU exceed 100,000. Most cancer possess 10 to 1000 deletions and insertions, but THCA-SA and OV-AU only carry about 10 deletions and insertions. Seven WGS projects identified multiple base substitutions. Five of them carry about 10 multiple base substitutions. ORCA-IN possesses more than 10 multiple base substitutions. In MELA-AU, numbers of multiple base substitutions range from 10 to more than 1000.

In Figure 1b, 10 to 1,000 single base substitutions were detected in the vast majority of WXS projects. Samples of LUCS-KR have the largest median (774) of single base substitutions. In CMDI-US and NBL-US, few single base substitutions were identified. WXS detected 10 times more deletions than insertions. The number of deletions in some cancers can exceed 1,000 but the maximum of insertion is just around 100. Multiple base substitutions were detected in six WXS projects. In four of them, there is only one multiple base substitution. In BRCA-UK and COCA-CN, the number increases to 10 and more.

### Spectra of SSM

It is clear in Figure 2 that C > T substitutions have the highest opportunity to occur. C > A and T > C are two other substitutions with high prevalence. The vast majority of indels are 1 to 3 bases in length. Multiple base substitutions only present in a few cancers. Multiple base substitutions of <= 20 bases are more prevalent than longer ones.

**Figure 2.**
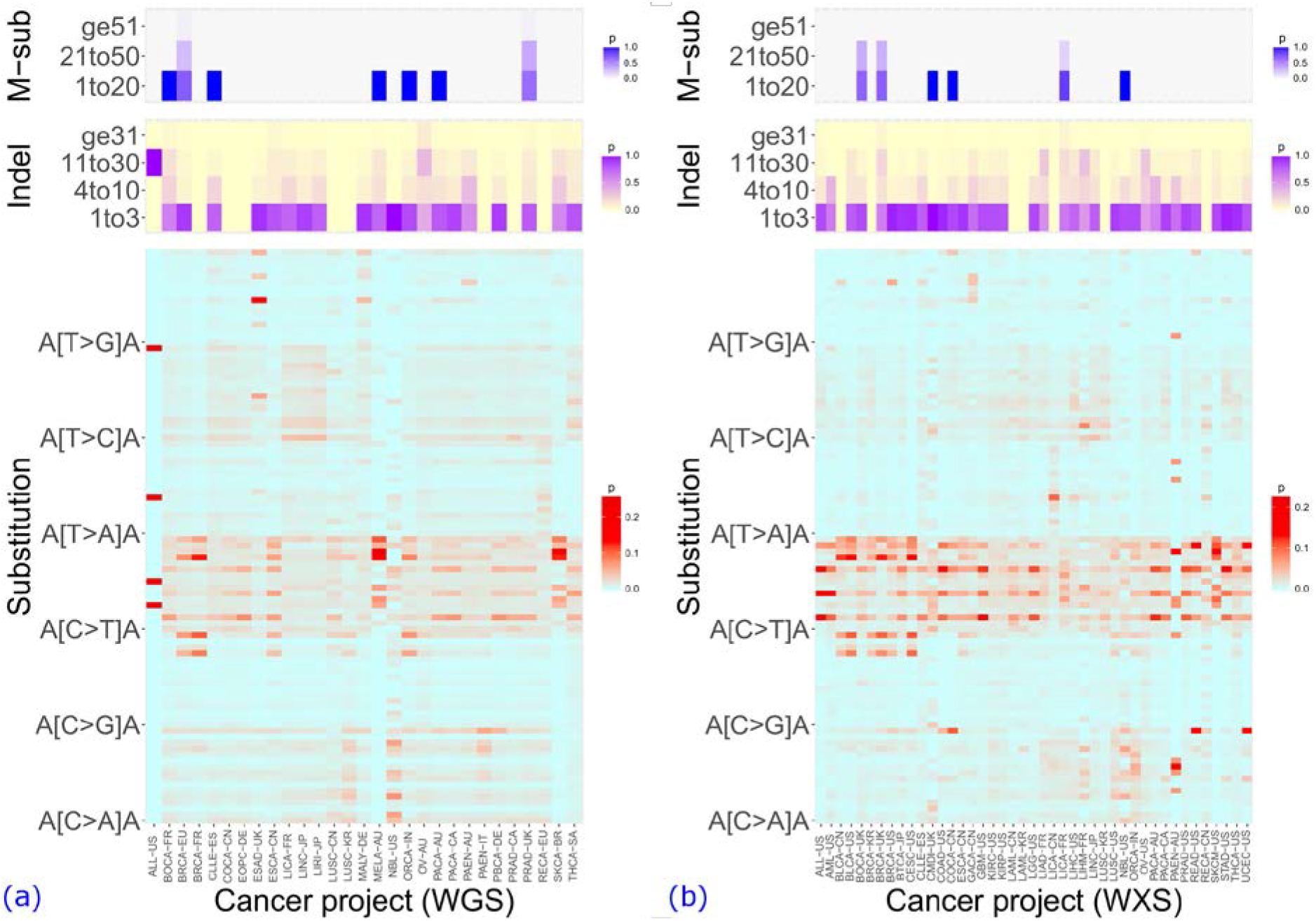
Spectra of SSM of 60 ICGC cancer projects. Substitution represents single base substitutions. There are 96 trinucleotide spectra for single base substitutions, in order of A[C>A]A/C/G/T, C[C>A]A/C/G/T, …, T[T>G]A/C/G/T, from bottom to top on y axis of the bottom panels. Indel are deletions and insertions. Indels are categorised as 1 to 3 bases (1to3), 4 to 10 bases (4to10), 11 to 30 bases (11to30), and greater than or equal to 31 bases (ge31). M-sub is multiple base substitutions. Spectra for M-sub are 1 to 20 bases (1to20), 21 to 50 bases (21to50), and greater than or equal to 51 bases (ge51). The heat map shows the percentage a spectrum takes in each category of SSM across cancer projects. (a) Cancer projects of WGS. (b) Cancer projects of WXS.

In WGS projects (Figure 2a), few C > A single base substitutions happen at NpCpG trinucleotides (N can be any base) but high prevalence of C > A transversions evenly distribute at other trinucleotides. BRCA-EU and BRCA-FR display similar spectra of substitution, with high probability of C > G and C > T at TpCpA and TpCpT. Unlike other cancers, ESAD-UK mainly has T > G at CpTpT. Two skin cancers, MELA-AU and SKCA-BR have almost the same spectra which are distinct from other cancers.Single base substitutions in the two skin cancers are enriched with C > T transitions that mainly occur at CpCpN (no CpCpG) and TpCpN.

There is an enrichment of C > T substitutions in WXS projects (Figure 2b). The C > T substitutions prefer to happen at NpCpG trinucleotides. Apart from high prevalence of C > T substitutions, BLCA-CN, BLCA-US, BRCA-KR, BRCA-UK, BRCA-US, and CESC-US similarly show high probability of C > G at TpCpA and TpCpT. COCA-CN, READ-US, and UCEC-US additionally have high prevalence of C > A at TpCpT. Less C > T single base substitutions appear in LICA-CN, but it is observed to have dominant T > A at CpTpG trinucleotide.

### Substitutions in chromosomes

Figure 3a demonstrates that all WGS cancers show similar distribution of single base substitutions per Mb across the chromosomes, with chromosome 8 per Mb having the most number of mutations and chromosome Y per Mb having the least. It is also clear to see that MELA-AU has an extremely high density of substitutions in every chromosome per Mb. It has at least three times more substitutions than other cancers per Mb.

**Figure 3.**
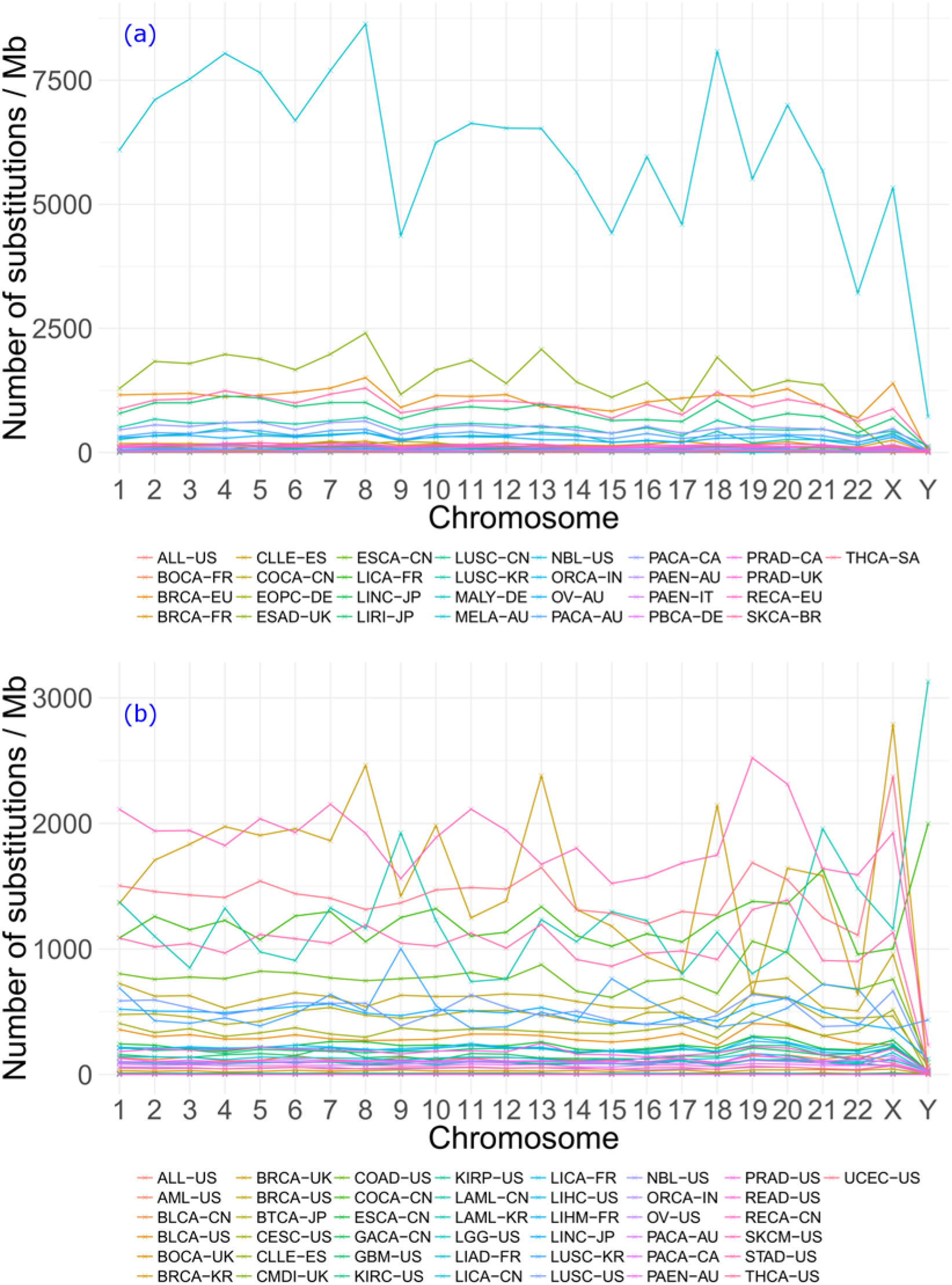
Distribution of number of single base substitutions per Mb across 24 chromosomes. (a) WGS projects. Chromosome size of human reference genome hg19 was applied to calculate the number of substitutions per Mb in each chromosome. (b) WXS projects. Size of exons on each chromosome was used to calculate the number of substitutions per Mb.

The interlacing lines in Figure 3b show no cancer can completely overtake others in terms of the distribution of single base substitutions per Mb per chromosome. In many WXS cancers, there is a peak in chromosome X, no matter how many substitutions per Mb in other chromosomes. COCA-CN, LAML-KR, and LUSC-KR have an obvious increase of substitutions in chromosome Y. In LAML-KR, the number stands above 3000 and exceeds other figures.

### Indels in chromosomes

According to Figure 4, the total number of indels in each chromosome varies in cancer projects, but it is noticeable that all projects display similar distribution of indels across chromosomes. In WGS projects (Figure 4a), there is a significant peak of number of indels in chromosome X. In WXS projects (Figure 4b), a peak of number of indels present in chromosome X as well. What’s more, there two other remarkable peaks in chromosome 17 and 19 in WXS cancer projects.

**Figure 4.**
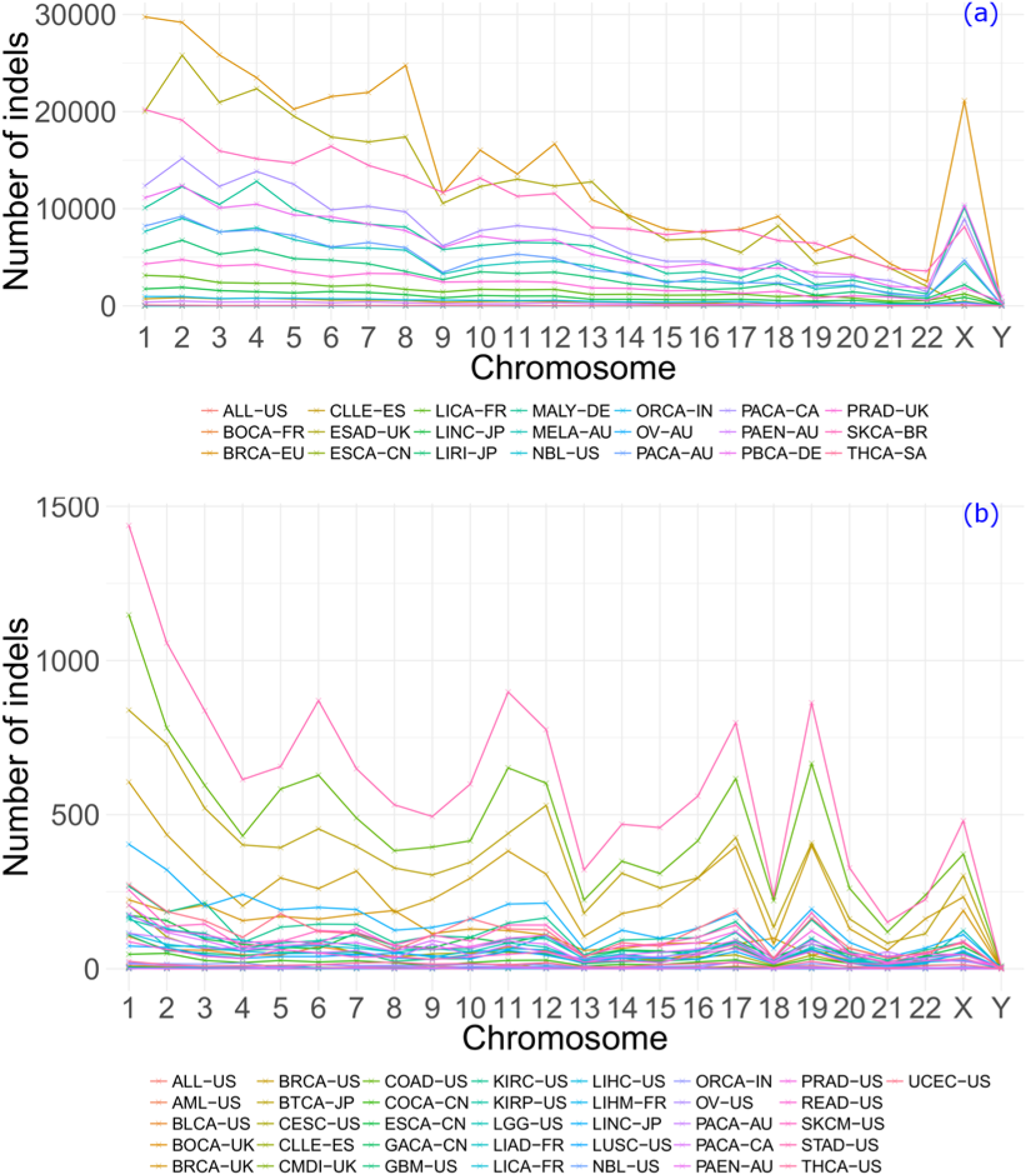
Distribution of total number of indels in 24 chromosomes. (a) Cancer projects of WGS. (b) Cancer projects of WXS.

## Discussion

There are only two samples in ALL-US sequenced by WGS. One sample carries four single base substitutions and the other has one deletion. The PAEN-AU in WXS projects only includes three samples, which contribute 14 single base substitutions, 16 deletions, and 6 insertions. The small number of mutations doesn’t bear statistical significance. Therefore, the results of WGS ALL-US and WXS PAEN-AU were only presented in the figures but not discussed in the context.

Compared to WXS, WGS detected 10 times more single base substitutions on average (Figure 1). The protein-coding regions constitute about 1% of human genome [16]. It is expected that more mutations should be called from the whole genome. As the ratio of length between genome and exome is not proportional to the ratio between numbers of substitutions, it is inferred that there is higher density of single base substitutions in the exome. This inference may also apply to deletions. WGS detected the same range of numbers of deletions and insertions. WXS detected 10 times more deletions than insertions. This may result from two causes: more insertions happen outside exons or insertions are more difficult to call [17]. In cancer projects which show multiple base substitutions, this mutation type was detected in almost all samples by WGS. WXS only detected multiple base substitutions in a small part of samples. To study this mutation type, the WGS strategy should be used with priority.

BRCA-EU (ER+ve, HER2-ve) and BRCA-FR (subtype defined by HER2) contains different number of WGS samples (560 vs. 64), but they have nearly the same spectra of single base substitutions. Their spectra are also very similar to another breast cancer, BRCA-UK (triple negative/lobular/other, 117 samples) in WXS projects. Their spectra are characterised by C > G and C > T at TpCpA and TpCpT. In two other breast cancers, BRCA-KR (Asian phenotype, 33 samples) and BRCA-US (ductal & lobular, 969 samples) of WXS, their spectra are characterised by C > G at TpCpA and TpCpT, and C > T at NpCpG. This shows that European breast cancers exhibit subtle difference from those outside Europe. This may unveil the difference between histologically different subtypes of breast cancer [18]. On the other hand, the high prevalence of C > G and C > T at TpCpA and TpCpT may widely exist in all breast cancers.

The most interest finding is that no prevalence of single base substitutions at NpCpG found in WGS BRCA-EU and WXS BRCA-UK. This preference was reported by the original publications of the samples in the two cancers [14,19]. Indeed, ours finding is not contradictory to the original one. In their articles, the fraction of 96 spectra of single base substitutions were normalized according to the prevalence of each trinucleotide in the genome. We didn’t normalize it that way. The frequencies of the 96 spectra were calculated as probabilities showing how likely 1 of the 96 substitutions can happen in the cancer. Our method not only shows the truth already known but also discloses the differences ignored previously.

It is also interesting to find similar spectra of single base substitutions between breast cancer (BRCA-EU, BRCA-FR, BRCA-KR, and BRCA-US), bladder cancer (BLCA-CN, BLCA-US), and cervical cancer (CESC-US). Breast cancer and cervical cancer are gender related. More than 90% cervical cancers are caused by human papillomavirus and bladder cancer has also found to be associated with viral infection [20,21]. The finding indicates it is worthwhile to study the common mechanisms underlying these cancers.

Different genomic characterisation exists between esophageal adenocarcinomas and esophageal squamous cell carcinomas [4]. This is evident in Figure 2a. Esophageal adenocarcinoma (ESAD-UK) and esophageal squamous carcinoma (ESCA-CN) show different spectra of single base substitutions. T > G transversions at CpTpT dominate single base substitutions in ESAD-UK whereas in ESCA-CN most substitutions are C > G at TpCpA and TpCpT and C > T at NpCpG.

WGS projects MELA-AU, SKCA-BR, and WXS SKCM-US are three skin cancers. The subtypes of MELA-AU and SKCM-US are melanoma and cutaneous melanoma. SKCA-BR is from skin and the type is melanoma, but the subtype was not given. Though from different countries, the three skin cancers have the same spectra of single base substitutions (Figure 2). Their distinct spectra from other cancers indicate the cancer originates from an absolutely specific aetiology. This may be attributed to the important role of exogenous carcinogens, mainly environmental factors like ultra violet, in the development of skin cancer [22,23].

In WGS projects, BRCA-EU and PRAD-UK has multiple base substitutions of more than 50 bases. No multiple base substitutions exceed 50 bases in WXS cancer projects. Multiple base substitutions were only detected in a few projects. This indicates this type of somatic mutations has not been well called and studied. Though it is hard to say which method is better at this moment, more studies are worthy of carrying out to study multiple base substitutions.

After excluding ALL-US, the distribution of single base substitutions per Mb of all WGS projects across chromosomes is drawn from the distribution of chromosome lengths (Table S3). In WXS projects, CDMI-UK, LAML-KR, LUSC-KR, NBL-US, and PAEN-AU don’t match the distribution of exons lengths across the chromosomes. This trend is also reflected by interlacing lines in Figure 3b. The different distribution of LAML-KR and LUSC-KR may result from high density of single base substitutions per Mb in chromosome Y. COCA-CN also has a larger number of substitutions per Mb in chromosome Y but doesn’t affect the distribution. The large number of substitutions of COCA-CN and LUSC-KR in chromosome Y may reflect the high risk of these cancers in men who have bad behaviours such as smoking and drinking [24,25]. The reason for many substitutions in chromosome Y of LAML-KR is unclear yet but may implicate certain mutation mechanisms of this cancer in men in South Korea.

## Conclusions

In this study, the pan-cancer scale landscape of SSM in 10878 samples of 60 ICGC cancer projects was offered. Many previous work towards mutational landscape only considered single base substitutions. Such limitation was overtaken in this study. Indels and multiple base substitutions were analysed, thus providing a more comprehensive landscape of SSM in human cancer. The difference between WGS and WXS strategies in cancer genome studies was compared according to the landscape of mutation at the pan-cancer scale the first time. Through the mutational landscape of SSM, not only knowledge already known was identified, novel mutational spectra in some cancers are revealed as well. Our study will enhance people’s understanding of somatic mutations in cancer genome. It can also help to study the mechanisms of somatic mutations underlying the development of cancer.

## Acknowledgements

This work is partly supported by National Natural Science Foundation of China (81373311, 31300674), China Scholarship Council (201506240135), and International Macquarie University Research Excellence Scholarships (2015233).

## Additional files

Additional file 1: Table S1, includes details of data used in this study, like cancer projects, cancer types, number of simple somatic mutations, number of samples and sequencing strategy. This is an Excel spreadsheet file (12KB)

Additional file 2: Table S2, includes the size of chromosomes for WGS and WXS. This is an Excel spreadsheet file (12KB).

Additional file 3: Table S3, includes the K-S result that shows whether the distribution of single base substitutions per Mb across chromosomes is drawn from the distribution of chromosome size in the genome. This an Excel spreadsheet file (12KB).

Additional file 4: Includes the statistics of simple somatic mutations that created the figures in this article. This is a compressed file and files inside are text files (zip 554 KB).

